# Temporal cueing improves auditory perception with no evidence for attentional tradeoffs

**DOI:** 10.1101/2025.11.11.687452

**Authors:** Juneau Wang, Christopher Conroy, Rachel N. Denison

## Abstract

Temporal cues help us focus attention at just the right moment. Whereas visual temporal attention selectively boosts perception at attended times at a cost to other moments, it remains unclear whether auditory temporal attention operates under the same constraints. To address this question, we designed a two-target temporal cueing task where participants discriminated the frequency sweep direction of one of two successive auditory stimuli, separated by 250 ms, while temporal attention was directed to one or both time points. We found that temporal attention improved auditory discrimination only for the first time point and could then remain sustained across the interval, without evidence of attentional tradeoffs in perceptual discriminability across time. Further, manipulating auditory feature uncertainty changed overall performance but did not modulate the impact of temporal attention or yield tradeoffs. Finally, comparison to previous data from a closely matched visual task confirmed that while temporal attention enhanced perception in both vision and audition, only vision shows tradeoffs at this timescale. Altogether the results suggest that the temporal constraints on attention are modality-dependent.

## Introduction

Whether watching for a tennis opponent to hit the ball after tracking the first part of their serve or listening for a subway announcement just after a chime, we often rely on temporal cues to help us focus attention at just the right moment. Voluntary temporal attention is the goal-directed prioritization of sensory information at specific moments in time, which improves perception and behavior during dynamic events (Nobre & van Ede, 2018). Previous research suggests that temporal attention operates as a limited yet recoverable resource, creating tradeoffs in perceptual performance over time (Denison et al., 2021). This dynamic allocation of attention has been characterized in vision using two-target temporal cueing tasks (Denison, 2024; Denison et al., 2017, 2021, 2024; Duyar et al., 2024; Fernández et al., 2019; Jing et al., 2022; Palmieri et al., 2023; Zhu et al., 2024), in which participants’ performance improves when attending to predictable, task-relevant stimuli and incurs costs when not attending, relative to a neutral condition in which participants distribute their attention across time.

A key open question is whether the attentional tradeoffs observed in vision also generalize to the auditory domain. Addressing this question is critical for determining whether temporal attention is constrained by a central, modality-independent resource or reflects distinct, sensory-specific processes. Although prior research has not addressed the question of temporal attentional tradeoffs in different modalities, there is evidence both that temporal aspects of attention operate similarly across modalities and, conversely, that auditory attention follows fundamentally different principles than visual attention.

On the one hand, temporal attention and expectation can have similar effects across sensory modalities. In audition, as in vision, individuals can prioritize specific moments in time to enhance perceptual discriminability. For example, auditory temporal expectation improves detection accuracy for tones occurring at anticipated time points (Wright & Fitzgerald, 2004). In vision, temporal expectation similarly enhances perceptual discriminability: targets appearing at anticipated time points are detected or identified more accurately (Correa et al., 2005; see also Correa et al., 2004). More broadly, both auditory and visual temporal attention and expectation involve both anticipatory and post-target processes (Denison et al., 2024; Lange et al., 2003).

Further evidence for parallels across modalities comes from studies of the attentional blink, the transient deficit in detecting a second target that follows a first by 200-500 ms. Although originally characterized in vision (Martens & Wyble, 2010; Raymond et al., 1992), analogous effects occur in audition (Duncan et al., 1997; Shen & Alain, 2011; Soto-Faraco & Spence, 2002) at similar timescales, which has been taken as evidence for a central processing bottleneck (Arnell & Jolicœur, 1999). Furthermore, in both modalities, temporal cueing can reduce the blink, improving detection of successive targets (Martens & Johnson, 2005; Shen & Alain, 2011).

Despite these parallels, temporal aspects of auditory and visual attention also show differences. The auditory attentional blink can have different dynamics than the visual attentional blink (Arnell & Jolicœur, 1999; Horváth & Burgyán, 2011; Potter et al., 1998). In addition, periodic temporal expectation appears tuned to modality-specific rhythms, with audition favoring faster rhythms (∼1.4 Hz) and vision favoring slower rhythms (∼0.7 Hz) (Zalta et al., 2020). Moreover, in non-temporal tasks, attentional resources for low-level auditory and visual tasks have been shown to operate largely independently, with minimal cross-modal interference during dual-task performance (Alais et al., 2006). It is therefore possible that while both the auditory and visual systems support temporal attention, they may do so via modality-specific implementations optimized for distinct sensory demands.

Although previous studies have suggested both shared and distinct mechanisms for attention across modalities, they have not isolated voluntary, or goal-directed, temporal attention in the auditory domain in a way that enables direct comparison with visual findings. Following a conceptual distinction developed for spatial and feature-based attention, expectation refers to the predictability of stimuli regardless of task-relevance, whereas attention refers to the relevance of a stimulus for the participant’s goal (Denison, 2024; Summerfield & Egner, 2009). Prior auditory studies—whether using temporally predictable versus unpredictable targets (Wright & Fitzgerald, 2004), rhythmic entrainment (Zalta et al., 2020), or rapid serial sequences engaging working memory (Shen & Alain, 2011)—have manipulated temporal attention together with expectation or stimulus-driven processes, limiting their ability to isolate voluntary temporal attention itself. Consequently, it remains unclear (1) whether voluntary temporal attention enhances auditory perception when separated from expectation and (2) whether it operates with tradeoffs at unattended times.

We addressed these questions using a two-target temporal cueing task adapted from the visual domain (Denison et al., 2017), which isolates temporal attention from expectation and quantifies its benefits and costs relative to a neutral baseline. In this task, two successive targets are presented and participants are instructed to attend to the first, the second, or both. If auditory temporal attention operates as a limited resource across the stimulus interval, then attending to the task-relevant moment should improve performance relative to attempting to sustain attention across time, making it strategic to attend to the target time (Denison, 2024). Moreover, attending to one stimulus would leave fewer resources available to process the other stimulus. Specifically, prioritizing either target should improve discrimination at the attended time (benefits) while impairing discrimination at the unattended time (costs), a pattern that has been found for visual temporal attention (Denison et al., 2017, 2021). Thus, the limited resource hypothesis predicts that temporal attention should affect both targets, as has typically been found in vision (Denison et al., 2017, 2021; Jing et al., 2022; Palmieri & Carrasco, 2024; Zhu et al., 2024).

Alternatively, if auditory temporal attention does not operate as a limited resource across the stimulus interval, then there would be no need to selectively attend to one time point over another. One possible outcome in this case is no effect of temporal attention on either target, as has been found for vision when the inter-stimulus interval is long (e.g., 800 ms) (Denison et al., 2021). Another possible outcome is that temporal attention would affect one target but not the other, as attention allocated to one target would not require a reduction in processing resources for the other target. In other words, if auditory attention does not operate as a limited resource over the stimulus interval, there should not be perceptual tradeoffs across time with auditory temporal attention. We evaluated these predictions across three experiments.

Further, across two experiments, we also manipulated feature uncertainty—via the range of possible target frequencies—to determine whether attentional effects vary depending on how predictable the relevant acoustic features are from trial to trial. Higher feature uncertainty has been shown to impair discrimination performance in vision (Irons et al., 2012; Kim et al., 2019; Mestry et al., 2017; Michel & Geisler, 2011; Pelli, 1985; Vogels et al., 1988) but may have weaker effects in some auditory detection tasks (Green, 1961; Tanner Jr., 1961). In a visual temporal attention task, feature uncertainty modulated the effects of temporal attention (Palmieri & Carrasco, 2024); thus, even if feature uncertainty has relatively weak effects on auditory performance itself, it may nevertheless interact with how temporal attention benefits auditory perception.

We found that temporal attention improves auditory discrimination only for the first time point and then could remain sustained without perceptual tradeoffs—a departure from prior findings in vision. Further, this attentional pattern was robust across levels of feature uncertainty. Finally, auditory performance gains were more modest than those previously observed for vision and exhibited a qualitatively different pattern of attentional benefits and costs across time, despite using a closely matched task across modalities, a pattern that provides further evidence against the limited resource hypothesis for auditory attention at this timescale. These findings reveal that voluntary temporal attention improves auditory perception, but apparently without the same temporal constraints found in vision, suggesting that resource limitations across time are not modality-general.

## Experiment 1

### Methods

#### Participants

Ten participants (7 females, ages 20–34, based on self-report) including author J.W. participated in Experiment 1. All participants except author J.W. were naive to the study. The sample size was selected to be the same as a previous study of visual temporal attention (Denison et al., 2017). All participants had normal hearing and normal or corrected-to-normal vision, based on self-report. All participants provided informed consent and were monetarily compensated, and the Boston University Institutional Review Board Involving Human Subjects approved the experimental protocols.

#### Stimuli

Stimuli were generated on a Linux computer using MATLAB 9.13 and Psychophysics Toolbox 3.0.17 (Brainard, 1997; Pelli, 1997; Kleiner et al., 2007). Visual stimuli were displayed on a gamma-corrected VPixx VIEWPixx monitor (VPixx Technologies Inc., QC, Canada) with a refresh rate of 120 Hz. Two monitors of the same model were used in the course of the study, with white luminance values of 92.9 and 97.1 cd/m^2^, respectively, and stimuli were presented on a 50% gray background. Auditory stimuli were presented through a Focusrite 2i2 3rd Gen audio interface (Focusrite, High Wycombe, Buckinghamshire, UK) in both ears with Sennheiser HD 569 headphones (Sennheiser, Lower Saxony, Wedemark, DE) with a sampling rate of 44100 Hz at a comfortable volume. Stimulus timing and audiovisual synchrony were verified using a photodiode and a PicoScope 2204A digital oscilloscope (Pico Technology, Cambridgeshire, UK). Participants’ heads were stabilized by a chin-and-head rest.

A central white fixation circle subtended 0.7°. Visual cues (30 ms) were given using two concentric rings (0.2° line width, diameter 2.0° and 2.9°) centered at fixation, where each ring could be either a darker 40% gray or white (**Figure 1**), similar to those used in previous temporal cueing studies (Coull et al., 2013, 2016; Capizzi et al., 2023). The ring colors served as temporal cues, with the white ring indicating which time point (inner: T1; outer: T2; both: neutral) should be attended or reported.

**Figure 1.**
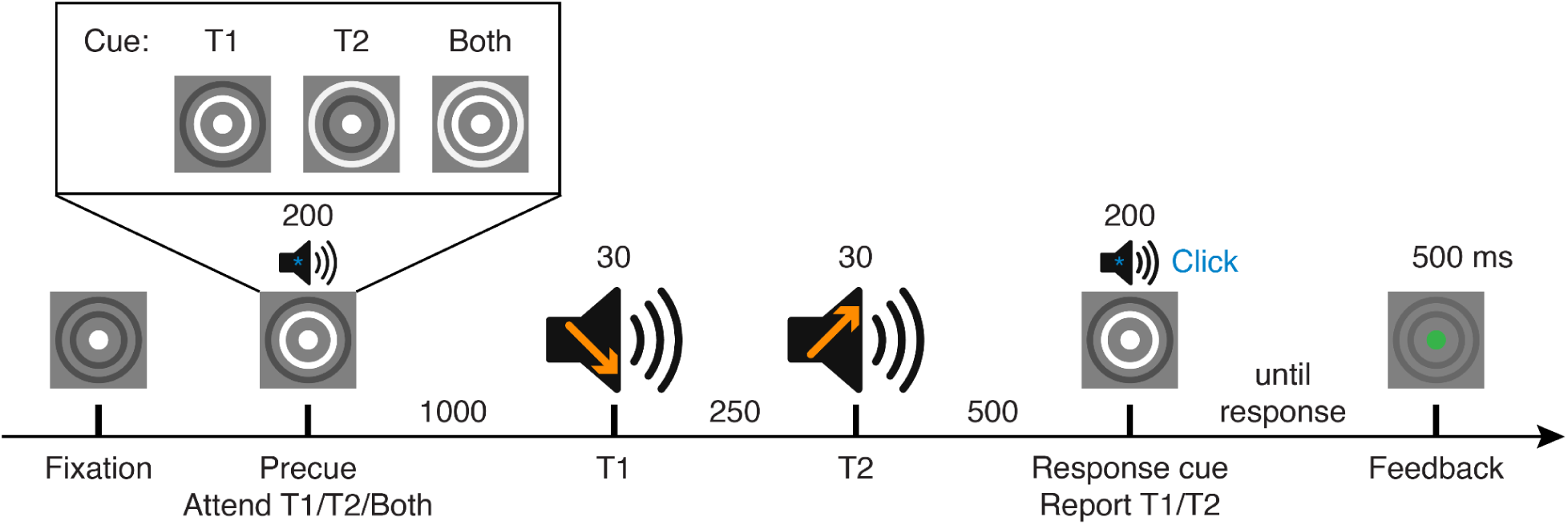
Auditory discrimination task. Participants discriminated the frequency sweep direction (up or down) of one of two sequential auditory stimuli, indicated by the response cue. A precue instructed the participant to attend to the first, second, or both targets. The trial timeline is well-matched to a previous visual temporal attention task (Denison et al., 2017). Timings give stimulus durations and SOAs.

The auditory cue was a brief (0.5 ms) “click,” a broadband noise ranging from 2–20 kHz. Auditory target stimuli were auditory sweeps that rapidly rose or fell in frequency over a 30 ms duration around a central frequency (**Figure 1**). For example, with a slope proportion of 0.3, the sweep ranged between ±30% of the central frequency (i.e., from 70% to 130% of the central frequency); so an 800 Hz upsweep therefore rose logarithmically from 560 Hz to 1040 Hz. Experiment 1 employed five central frequencies that were logarithmically-spaced from 800–2400 Hz. A single threshold slope value was estimated and used across all frequencies. Because the slope was proportional to the central frequency, sweeps for different central frequencies spanned different ranges but their magnitudes were equated on a log scale. All auditory stimuli were enveloped with cosine-squared ramps (targets, 5 ms wide; cues, 0.25 ms wide) to smooth the onset and offset.

#### Procedure

Participants performed a two-target temporal cueing task while directing attention to one of two auditory frequency sweeps played in sequence (**Figure 1**). The task was designed as an auditory version of the visual temporal cueing task introduced by Denison et al. (2017). On each trial, two auditory frequency sweeps (targets T1 and T2), with independent sweep directions (up or down) and central frequencies, were presented for 30 ms each and separated by a 250-ms stimulus onset asynchrony (SOA). The timing of each trial was predictable. We used a frequency sweep discrimination task because determining the direction of an auditory sweep shares properties with discriminating the tilt of a visual line or grating. This stimulus also allowed task difficulty to be titrated for each participant. Multiple central frequencies were employed to minimize trials in which the two stimuli were identical, discouraging participants from relying on a similarity judgment strategy.

A visual precue ring, presented 1000 ms before T1, instructed participants to attend to either the first or the second target in selective attention trials (80% of trials, inner ring: T1; outer ring: T2) or to sustain attention across both targets in neutral trials (20% of trials, both rings presented simultaneously). Visual precues were used to avoid presenting any auditory stimuli with specific frequency content that could bias subsequent frequency sweep processing. A broadband auditory click was presented together with the visual precue to ensure the participant had precise precue timing information. A visual response cue ring, presented 500 ms after T2, instructed participants to report the frequency sweep direction (up or down) of a single target (inner ring: T1; outer ring: T2). The response cue matched the precue with 75% validity in selective attention trials and was equally likely to indicate each target in neutral trials.

Participants reported the direction (up: key 1, down: key 2) of the auditory sweep indicated by the response cue. Participants received feedback via a change in fixation color (500 ms, correct: green; incorrect: red) after each trial and performance accuracy feedback (percent correct) after each block of 100 trials.

Participants completed two 75-minute sessions on separate days. A total of 1,000 trials represented all counterbalanced combinations of cue type (valid: 60%, neutral: 20%, invalid: 20%), target (T1, T2), target sweep direction (up, down; independent for T1 and T2), and target central frequency (800 Hz, 1053 Hz, 1386 Hz, 1824 Hz, 2400 Hz; independent for T1 and T2) in a randomly shuffled order. These trials were divided across two sessions as 10 blocks of 100 trials, with 20 or more practice trials at the start of each session.

#### Training

Participants completed one or two 75-minute sessions of training before the experiment to familiarize them with the task and determine their sweep slope thresholds. Each training step except thresholding used blocks of 20 trials, which were repeated until the participant reached 75% accuracy. The training steps were as follows: (1) Slow, easy, neutral: 1,000 ms SOA, all neutral precues, and a sweep slope ranging from 0.3–0.6 proportion of the central frequency based on what was easily discriminable for the participant. (2) Fast, easy, neutral: 250 ms SOA, all neutral precues, and an easily discriminable sweep slope. (3) Thresholding: 160 trials with two interleaved 3-up-1-down staircases (one starting easy, one starting difficult) with all neutral precues but otherwise identical to the main task to determine participants’ individual sweep slope thresholds. These sweep slope values were used for subsequent training steps. (4) Main task, valid: main task but with all valid precues. (5) Main task: all precue types, identical to experimental trials.

Across days, participants began each experimental session by repeating steps 4–5 as a warm-up. If performance during this warm-up did not reach 75% accuracy, step 3 (thresholding) was repeated and the newly estimated sweep slope was used for the experimental blocks that followed.

#### Analysis

To assess the effects of temporal attention on perceptual discriminability, we used discrimination accuracy as the dependent measure. Reaction time was collected as a secondary measure and was used to confirm that participants were following task instructions.

We used generalized and linear mixed-effects models to analyze accuracy and reaction time (RT) data across three experiments. All models were followed by Type II Wald chi-square tests using the Anova() function from the *car* package in R.

We analyzed trial-level accuracy (binary outcome, correct or incorrect) using binomial generalized linear mixed-effects models (GLMMs) with a logit link. Models of all trials included fixed effects of validity (valid, neutral, invalid) and target (T1, T2) with subject-specific random intercepts. When responses to each target were analyzed separately, models included only validity as a fixed effect and subject-specific random intercepts. Target-specific analyses were planned based on the theoretical expectation that T1 and T2 may exhibit different patterns due to the inherent directionality of time. Pairwise comparisons of validity (valid vs. invalid, valid vs. neutral, and neutral vs. invalid) were planned to separately assess attentional benefits (valid > neutral), costs (invalid < neutral), and overall effects (valid > invalid). Each contrast was tested using separate two-level GLMMs restricted to the relevant conditions, following the same model structure described above.

We analyzed trial-level RTs using linear mixed-effects models (LMMs). To correct for positive skew and approximate normality, we first log-transformed the RTs. This transformation was selected using model comparison: among candidate distributions (Gaussian, log-normal, inverse Gaussian, Gamma), the log-normal distribution yielded the lowest average AIC and BIC values across subjects. The RT models included the same fixed and random effects structure as the accuracy models. Because this is an unspeeded task, we chose a conservative criterion for trial exclusion: two trials with RTs exceeding 60 seconds were excluded, which was equivalent here to excluding all trials with RTs exceeding 3 standard deviations above the mean in a log-normal RT distribution; their removal did not affect any statistical outcomes in either accuracy or RT analyses.

## Results

On each trial, participants heard two sequential auditory frequency sweeps (targets, T1 and T2) and were instructed using a precue to attend to one of the two targets, or both targets in the neutral condition. Participants judged the direction of the frequency sweep (up or down) for the target indicated by a response cue at the end of the trial. Critically, the precued and response-cued targets matched on most trials but not always, allowing us to compare performance across valid, neutral, and invalid precueing conditions. The timing of the two targets was fully predictable and only the timing of attention varied from trial to trial with the precue, allowing us to isolate the effects of temporal attention on auditory discrimination accuracy and RT. We assessed whether voluntary temporal attention could improve perceptual discriminability to auditory stimuli (valid vs. invalid) and whether any such improvements occurred for T1, T2, or both targets. Further, we assessed whether attending to one target would improve perception beyond attending to both, and whether any such attentional benefits (valid vs. neutral) would be accompanied by perceptual impairments (costs, neutral vs. invalid) at unattended time points, providing evidence for attentional selectivity across time.

### Temporal attention improved auditory discriminability for T1 but not T2

Temporal attention modestly enhanced discrimination of auditory sweep direction for T1 (**Figure 2a**), with accuracy higher for valid and neutral trials than for invalid trials—a trend that approached significance (*χ²*(2) = 5.53, *p* = 0.063). This pattern reflects a cost of inattention, with significantly lower accuracy for invalid than valid trials (*χ²*(1) = 4.47, *p* = 0.035) and for invalid compared to neutral trials (*χ²*(1) = 4.30, *p* = 0.038). However, there was no significant difference between valid and neutral trials (*χ²*(1) = 0.21, *p* = 0.649). Thus attending to T2 produced a performance cost to T1, but attending to T1 produced no benefit over attending to both targets.

**Figure 2.**
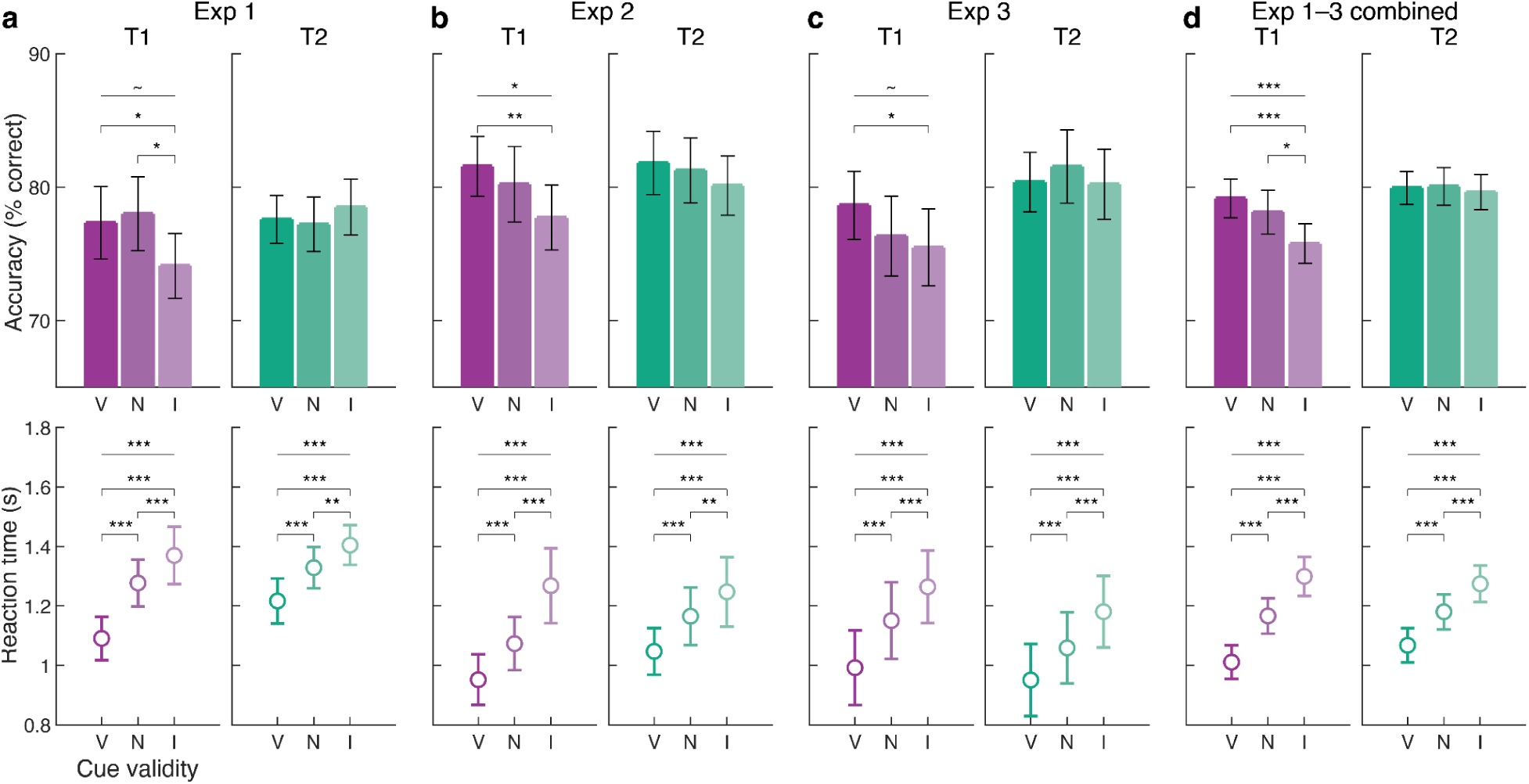
Mean accuracy and reaction time for each temporal cueing condition in **(a)** Experiment 1 (n = 10 participants), **(b)** Experiment 2 (n = 10), **(c)** Experiment 3 (n = 11), and **(d)** all experiments combined. Horizontal lines show main effects of validity within a target, while brackets show pairwise comparisons between precue conditions. Error bars represent ±1 SEM. V = valid, N = neutral, I = invalid. ∼ p < 0.1, * p < 0.05, ** p < 0.01, *** p < 0.001.

For T2, temporal attention had no effect on sweep direction discrimination. There was no significant main effect of validity (*χ²*(2) = 0.55, *p* = 0.759), nor any significant pairwise difference between valid, neutral, or invalid trials (valid vs invalid: *χ²*(1) = 0.38, *p* = 0.535; valid vs neutral: *χ²*(1) = 0.06, *p* = 0.809; neutral vs invalid: *χ²*(1) = 0.50, *p* = 0.480). Thus auditory temporal attention improved T1 discriminability (valid vs. invalid) without any apparent tradeoffs for T2. That is, effects of attention for T1 were not accompanied by corresponding attentional effects for T2.

The different patterns of attentional effects for T1 and T2 were reflected in a trending interaction between validity and target (*χ²*(2) = 4.62, *p* = 0.099). There was no main effect of validity on auditory discriminability across the two targets (*χ²*(2) = 1.40, *p* = 0.495), likely driven by the absence of T2 effects. Overall auditory discriminability was not significantly different for the two targets (main effect of target, *χ*²(1) = 1.09, *p* = 0.297), indicating that the lack of attention effects for T2 were not tied to overall performance differences.

### Temporal attention improved reaction time for both targets

Reaction time was strongly modulated by the temporal precue (**Figure 2a**), with faster responses for valid trials compared to neutral trials, which were in turn faster than invalid trials across both T1 and T2 (main effect of validity, *χ²*(2) = 346.52, *p* < 1e-15). This pattern of attention was consistent for each target separately. For T1, temporal attention reduced RTs (main effect of cue validity, *χ²*(2) = 251.03, *p* < 1e-15), with all pairwise comparisons showing reliable differences (valid < neutral: *χ²*(1) = 103.45, *p* < 1e-15; neutral < invalid: *χ²*(1) = 11.55, *p* = 6.79e-4; valid < invalid: *χ²*(1) = 204.17, *p* < 1e-15). The same was true for T2, with a significant main effect (*χ²*(2) = 112.64, *p* < 1e-15) and all pairwise contrasts significant (valid < neutral: *χ²*(1) = 38.34, *p* = 5.96e-10; neutral < invalid: *χ²*(1) = 9.37, *p* = 0.002; valid < invalid: *χ²*(1) = 97.16, *p* < 1e-15). T1 showed overall faster RTs compared to T2 (main effect of target, *χ²*(1) = 84.04, *p* < 1e-15) along with more pronounced attentional effects (validity by target interaction, *χ²*(1) = 13.62, *p* = 0.001). These results confirm that participants were following the task instructions for both targets and that the observed effects of attention on perceptual discriminability cannot be explained by a speed–accuracy tradeoff (Green & Luce, 1972).

## Experiment 2

We next asked whether and how auditory temporal attention interacted with feature uncertainty, or the predictability of stimulus features such as auditory frequency or visual orientation. Experiment 2 reduced feature uncertainty compared to Experiment 1 by limiting the number of central frequencies to two, rather than five, thereby making the stimulus frequencies more predictable, which we expected would reduce task difficulty arising from feature uncertainty while still titrating the frequency sweep slope to equate performance with Experiment 1 (**Figure 3a**). This design not only decreases the range of auditory features that participants need to encode and compare, but also aligns more closely with the visual version of the two-target temporal cueing task, in which stimuli varied around two feature values (i.e., vertical or horizontal orientations) (Denison et al., 2017). By simplifying the auditory stimulus space, we aimed to test whether reducing feature uncertainty would lead to stronger effects of temporal attention on perceptual discriminability compared to Experiment 1.

**Figure 3.**
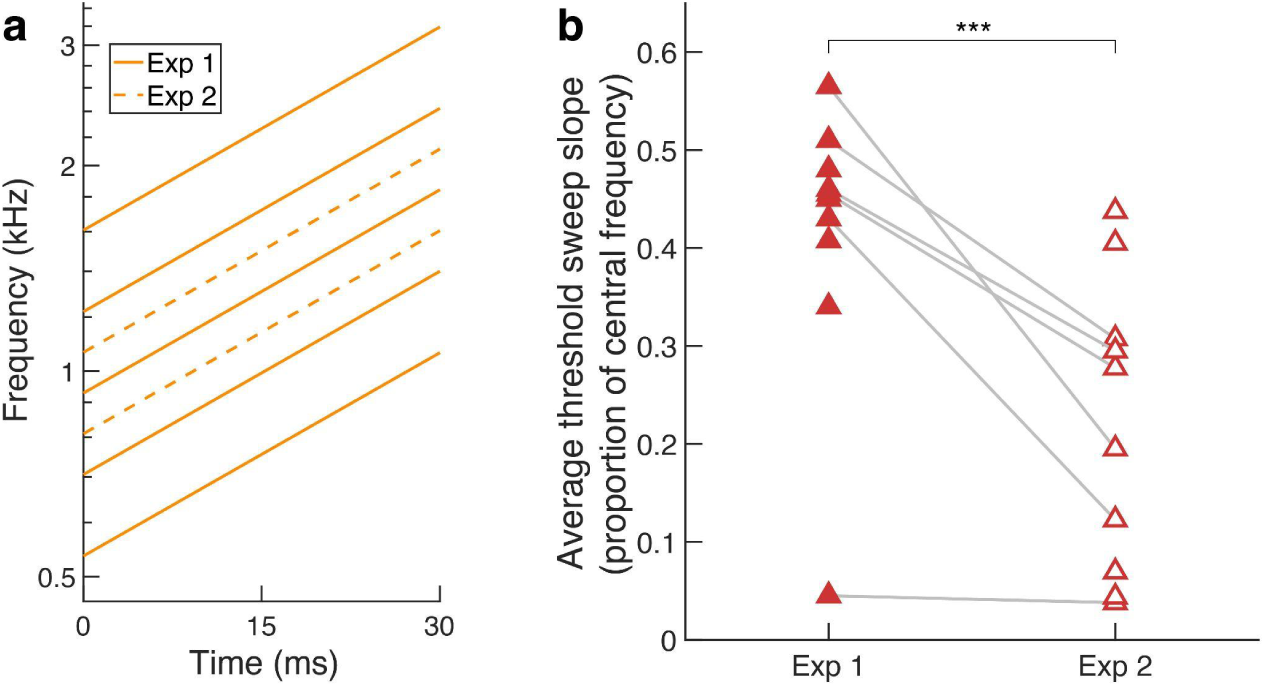
Reducing the number of central frequencies improved thresholds. **(a)** Example upward auditory sweep stimuli with a slope proportion of 0.33 of the central frequency. The y-axis is logarithmic. **(b)** Threshold sweep slope averaged across sessions for each participant in Experiments 1 and 2. n = 10 participants per experiment. *** p < 0.001.

## Methods

In Experiment 2, the methods were the same as in Experiment 1 except for the differences described below.

### Participants

Ten participants (6 females, ages 18–35, based on self-report) participated in Experiment 2. Author J.W. and five other participants who participated in Experiment 1 also participated in Experiment 2.

### Stimuli

Experiment 2 used two central frequencies (1208 Hz and 1590 Hz) rather than five central frequencies. These central frequencies were selected to fall within the Experiment 1 range and still be highly discriminable.

### Procedure

A total of 960 trials represented six repetitions of all combinations of cue type (valid: 60%, neutral: 20%, invalid: 20%), target (T1, T2), target sweep direction (up, down; independent for T1 and T2), and target central frequency (1208 Hz, 1590 Hz; independent for T1 and T2) in a randomly shuffled order. Trials were divided into two sessions as 6 blocks of 160 trials, with 32 or more practice trials at the start of each session. Trial conditions were counterbalanced across each session.

### Training

The training procedure matched that of Experiment 1, except it used blocks of 32 trials and included two additional training steps at the end: (1) a full practice run of 160 trials with all precues, identical to experimental trials; (2) a second staircase of 160 trials to ensure that thresholds were stable.

### Analysis

To compare task difficulty between Experiments 1 and 2, we compared sweep slopes (the mean slope used across each participant’s experimental sessions) using an LMM.

In the trial-level data analysis, one trial was excluded due to an RT exceeding 60 seconds. Removing this trial did not affect any statistical outcomes in either accuracy or RT analyses.

## Results

### Task difficulty decreased with reduced feature uncertainty

Sweep discrimination thresholds in Experiment 2 were significantly lower than in Experiment 1 (*χ*²(1) = 19.70, *p* = 9.05e-06), demonstrating that the reduction in feature uncertainty reduced task difficulty (**Figure 3b**).

### Temporal attention improved auditory discriminability for T1 but not for T2

Temporal attention enhanced discrimination of auditory sweep direction for T1 (**Figure 2b**), with accuracy highest for valid, intermediate for neutral, and lowest for invalid trials (*χ²*(2) = 7.05, *p* = 0.029). Attention improved performance for valid relative to invalid trials (*χ²*(1) = 6.99, *p* = 0.008). These results confirm an effect of auditory temporal attention on T1. However, there was no significant difference between valid and neutral trials (*χ²*(1) = 0.88, *p* = 0.348) or invalid and neutral trials (*χ²*(1) = 1.87, *p* = 0.172).

As in Experiment 1, attention did not improve auditory discriminability for T2. There was no significant main effect of validity (*χ²*(2) = 1.42, *p* = 0.491), nor any significant pairwise difference between valid, neutral, or invalid trials (valid vs invalid: *χ²*(1) = 1.42, *p* = 0.234; valid vs neutral: *χ²*(1) = 0.15, *p* = 0.695; neutral vs invalid: *χ²*(1) = 0.42, *p* = 0.519).

Across both targets, attention enhanced auditory discriminability (*χ*²(2) = 7.43, *p* = 0.024). However, there was no main effect of target (*χ*²(1) = 1.11, *p* = 0.291) nor interaction between validity and target (*χ²*(2) = 0.98, *p* = 0.612). Together, these results indicate that reducing feature uncertainty effectively decreased task difficulty but did not qualitatively alter the pattern of temporal attention effects across targets.

### Temporal attention improved reaction time for both targets

Reaction time again was strongly modulated by the temporal precue (**Figure 2b**), with faster responses for valid trials compared to neutral trials, which were in turn faster than invalid trials across both T1 and T2 (main effect of validity, *χ*²(2) = 323.03, *p* < 1e-15). This pattern of attention was consistent for each target separately. For T1, temporal attention reduced RTs (main effect of cue validity, *χ²*(2) = 241.93, *p* < 1e-15), with all pairwise comparisons showing reliable differences (valid < neutral: *χ²*(1) = 55.28, *p* = 1.05e-13; neutral < invalid: *χ²*(1) = 38.98, *p* = 4.28e-10; valid < invalid: *χ²*(1) = 228.62, *p* < 1e-15). The same was true for T2, with a significant main effect (*χ²*(2) = 103.36, *p* < 1e-15) and all pairwise comparisons significant (valid < neutral: *χ²*(1) = 37.68, *p* = 8.34e-10; neutral < invalid: *χ²*(1) = 7.26, *p* = 0.007; valid < invalid: *χ²*(1) = 88.08, *p* < 1e-15). As in Experiment 1, T1 also showed overall faster RTs compared to T2 (main effect of target, *χ*²(1) = 62.05, *p* = 3.35e-15) along with more pronounced attentional effects (validity by target interaction, *χ*²(2) = 13.23, *p* = 0.001).

Again, the improvements in RT with cue validity indicate that participants were performing the task correctly and that the effects of the precue on accuracy cannot be explained by a speed–accuracy tradeoff. Instead, the effects of the precue on accuracy reflect differences in perceptual processing.

## Testing the interaction of feature uncertainty and temporal attention

Although Experiment 2 had reduced central frequency uncertainty compared to Experiment 1, both experiments yielded qualitatively similar patterns of results—namely, an effect of auditory temporal attention on T1 but not T2 discrimination. We next directly compared Experiments 1 and 2 to assess whether feature uncertainty modulated the effects of temporal attention.

## Methods

### Analysis

We used the same trial-level modeling approach as in Experiments 1 and 2: binomial GLMMs for accuracy and log-transformed LMMs for RT. Models for the combined dataset also include experiment as a factor, allowing tests of whether temporal attention effects varied across feature uncertainty levels. If no significant interactions involving experiment were found across all models, we planned to evaluate main effects of validity and target across uncertainty levels using the combined data.

For critical nonsignificant pairwise comparisons, we additionally performed equivalence tests using the Two One-Sided Tests (TOST) procedure (Lakens, 2017) to test whether accuracy differences between any two validity conditions were smaller than 2.5%. We also quantified evidence for or against validity effects using Bayes factor (BF10). For each target, we compared a model including validity as a fixed factor (H1) with a reduced model excluding validity (H0) using *lmBF()* from the *BayesFactor* package in R (Rouder et al., 2017). As with the mixed-effects models, the Bayes factor model for the combined dataset included experiment as a fixed factor. Then, to quantify the evidence for or against any interaction of feature uncertainty and temporal attention, we compared a combined data model with target as an additional factor and including the interaction as a fixed factor (H1) with a reduced model excluding the interaction (H0). BF10 values greater than 1 indicate evidence in favor of an effect, whereas values less than 1 indicate evidence favoring the null model (no effect) (Kass & Raftery, 1995; Lee & Wagenmakers, 2014). The strength of evidence was interpreted according to the recommendations of Lee & Wagenmakers (2014, section 7.2).

## Results

### Feature uncertainty modulated task difficulty but not attentional effects

Performance improved with reduced feature uncertainty (main effect of experiment, accuracy: *χ²*(1) = 47.32, *p* = 6.02e-12; RT: *χ²*(1) = 364.78, *p* < 1e-15), with higher accuracy and lower RT for Experiment 2 compared to Experiment 1 (**Figure 2a,b**). Removing the highest and lowest central frequencies from Experiment 1 (800 Hz, 2400 Hz) may have contributed to this improvement, because they had lower performance overall compared to the more intermediate central frequencies. However, performance for both Experiment 2 central frequencies (1208, 1590 Hz) matched or exceeded performance at the easiest two Experiment 1 central frequencies (1053, 1386 Hz) for both accuracy (main effect of experiment: *χ²*(1) = 9.73, *p* = 0.002) and reaction time (main effect of experiment: *χ²*(1) = 219.43, *p* < 1e-15), even with lower sweep discrimination thresholds. This pattern supports the conclusion that reduced feature uncertainty made the task easier in Experiment 2.

However, critically, we found no interactions of experiment with validity or target (all *χ*² ≤ 2.98, df = 1–2, all *p*s ≥ 0.226) for either accuracy or RT. In the target-specific models, there were main effects of experiment (all *χ*²(1) ≥ 15.05, all *p*s ≤ 1.05e-4) but no interactions of experiment and validity (all *χ*²(2) ≤ 5.41, all *p*s ≥ 0.067) for either accuracy or RT. Furthermore, Bayes Factor analysis provides moderate evidence against an interaction between feature uncertainty and temporal attention (BF10 = 0.16). Therefore, although overall performance differed between experiments, the effect of temporal attention (validity) was stable across different levels of feature uncertainty.

## Experiment 3

We then replicated Experiment 2, but with a delayed response period. There were two motivations for conducting this experiment: (1) to generate a larger combined dataset with higher statistical power for assessing the effects of auditory temporal attention; and (2) to ensure any effects observed in RT are unlikely to be driven by changes in perceptual processing speed at the time of the stimulus itself, as the interval between the targets and response period was set to be 1.1 s, much longer than the expected duration of perceptual processing (Donkin et al., 2014; Vangkilde et al., 2012).

## Methods

In Experiment 3, the methods were the same as in Experiment 2 except for the differences described below.

### Participants

Eleven participants (4 females, ages 25–36, based on self-report) participated in Experiment 3. We recruited eleven participants aiming for a target number of ten participants to match the other experiments, given typical rates of participant drop-out in multi-session experiments, but all recruited participants completed the study.

### Stimuli

A central white fixation circle subtended 0.6°. Visual cues (30 ms) were given using numbers in black font within the fixation circle indicating which time point (“1”: T1; “2”: T2, “12”: neutral) should be attended (precue) or reported (response cue). The average luminance of the cues was approximately equated by adjusting the font size to match the number of black pixels across cues (“1”: 43 px; “2”: 47 px; “12”: 47 px). We used number cues instead of ring cues in this experiment to reduce the training needed to learn the cue mapping.

### Procedure

The procedure was identical to Experiment 2, except that participants could not respond until a “go” cue (dimming of the fixation circle to 40% gray) appeared 600 ms after the onset of the response cue **(Figure 4)**. Participants reported both the direction of the auditory sweep indicated by the response cue and their confidence in their judgment (up, high confidence: key 1; up, low confidence: key 2; down, low confidence: key 9; down, high confidence: key 0). Confidence results will not be reported in this manuscript. Fixation was monitored using an EyeLink 1000 Plus eye tracker (SR Research, Ottawa, Ontario, CA) with a sampling frequency of 1000 Hz.

**Figure 4.**
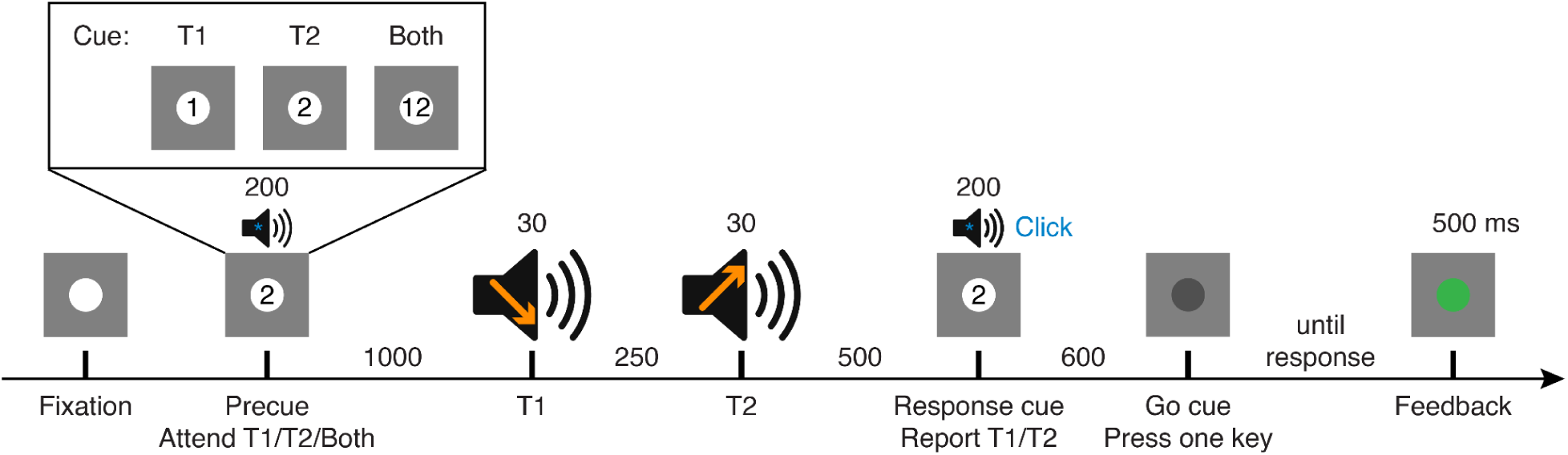
Auditory discrimination task for Experiment 3, which introduced a delay before participants were allowed to respond. Timings give stimulus durations and SOAs.

### Training

The training procedure matched that of Experiment 1, except it used blocks of 32 trials.

### Analysis

Reaction times were recorded from the onset of the “go” cue rather than the onset of the response cue.

## Results

### Temporal attention improved auditory discriminability for T1 but not for T2

Temporal attention again modestly enhanced discrimination of auditory sweep direction for T1 (**Figure 2c**). Accuracy was highest for valid, intermediate for neutral, and lowest for invalid trials—a trend that approached significance (*χ²*(2) = 5.86, *p* = 0.053). Attention improved performance for valid relative to invalid trials (*χ²*(1) = 4.74, *p* = 0.029). However, there was no significant difference between valid and neutral trials (*χ²*(1) = 2.55, *p* = 0.110) or invalid and neutral trials (*χ²*(1) = 0.22, *p* = 0.640). As in Experiments 1 and 2, attention did not improve auditory discriminability for T2. There was no significant main effect of validity (*χ²*(2) = 0.83, *p* = 0.660), nor any significant pairwise difference between valid, neutral, or invalid trials (valid vs invalid: *χ²*(1) = 0.01, *p* = 0.910; valid vs neutral: *χ²*(1) = 0.72, *p* = 0.397; neutral vs invalid: *χ²*(1) = 0.63, *p* = 0.428). Across both targets, there was a main effect of target (*χ*²(1) = 15.06, *p* = 1.04e-4) but no main effect of attention (*χ*²(2) = 2.75, *p* = 0.253) nor interaction between validity and target (*χ²*(2) = 3.91, *p* = 0.142). Thus, consistent with Experiments 1 and 2, temporal attention improved auditory discrimination for T1 but not for T2.

### Temporal attention improved reaction time for both targets

Reaction time again was strongly modulated by the temporal precue (**Figure 2b**), with faster responses for valid trials compared to neutral trials, which were in turn faster than invalid trials across both T1 and T2 (main effect of validity, *χ*²(2) = 341.76, *p* < 1e-15). This pattern of attention was consistent for each target separately. For T1, temporal attention reduced RTs (main effect of cue validity, *χ²*(2) = 195.56, *p* < 1e-15), with all pairwise comparisons showing reliable differences (valid < neutral: *χ²*(1) = 62.52, *p* = 2.64e-15; neutral < invalid: *χ²*(1) = 20.90, *p* = 4.85e-06; valid < invalid: *χ²*(1) = 162.79, *p* < 1e-15). The same was true for T2, with a significant main effect (*χ²*(2) = 150.69, *p* < 1e-15) and all pairwise comparisons significant (valid < neutral: *χ²*(1) = 42.67, *p* = 6.47e-11; neutral < invalid: *χ²*(1) = 21.00, *p* = 4.59e-06; valid < invalid: *χ²*(1) = 129.16, *p* < 1e-15). As in Experiments 1 and 2, T1 also showed overall faster RTs compared to T2 (main effect of target, *χ*²(1) = 13.00, *p* = 3.12e-4). The effects of the precue on T1 and T2 were similar (validity by target interaction, *χ*²(2) = 1.44, *p* = 0.486). Therefore, precue validity affected RT even with an extended (1.1 s) interval between the targets and the response window.

## Combined results

Given the comparable results in Experiments 1, 2, and 3, with no impact of feature uncertainty on the effects of temporal attention when comparing Experiments 1 and 2, we next combined the datasets from the three experiments to obtain more power to estimate the effects of temporal attention on auditory perception (**Figure 2d**).

### Temporal attention improved auditory discriminability for T1 but not for T2

The combined dataset showed an overall effect of attention on auditory discriminability (χ²(2) = 9.71, p = 0.008), with an interaction between validity and target indicating a greater effect of temporal attention on T1 than on T2 (*χ*²(2) = 6.28, *p* = 0.043). Temporal attention strongly improved T1 sweep discrimination (main effect of validity: *χ²*(2) = 15.86, *p* = 3.59e-4; valid vs. invalid: *χ²*(1) = 15.84, *p* = 6.90e-05). These improvements reflected costs of inattention: accuracy was lower for invalid compared to neutral trials (*χ²*(1) = 4.99, *p* = 0.026) but not significantly different between valid and neutral trials (*χ²*(1) = 1.47, *p* = 0.226). A TOST equivalence test confirmed that any accuracy difference between valid and neutral trials was significantly smaller than 2.5% (estimate: 0.99% ± 0.83% SE, TOST *p* = 0.034), indicating no detectable attentional benefits beyond the neutral baseline. Further, Bayes Factor analysis provides anecdotal evidence for no accuracy difference between valid and neutral trials for T1 (BF10 = 0.35).

Conversely, attention did not improve auditory discriminability for T2, with no significant main effect of validity (*χ*²(2) = 0.179, *p* = 0.915), nor any significant pairwise differences between valid, neutral, or invalid trials (valid vs invalid: *χ*²(1) = 0.128, *p* = 0.721; valid vs neutral: *χ*²(1) = 0.02, *p* = 0.899; neutral vs invalid: *χ*²(1) = 0.15, *p* = 0.695). All TOST equivalence tests confirmed that any pairwise differences were smaller than 2.5% accuracy (valid – invalid: 0.32% ± 0.81% SE; valid – neutral: 0.13% ± 0.80% SE; neutral – invalid: −0.45% ± 0.99% SE; all TOST *p*s ≤ 0.019). Bayes Factor analysis provides moderate evidence for no overall effect of validity on T2 (BF10 = 0.19).

Overall accuracy was significantly higher for T2 than for T1 (main effect of target: χ²(1) = 12.12, p = 5.00e-4), which was primarily driven by Experiment 3 (**Figure 2c,d**). Meanwhile, as expected from the individual experiment results, RT improved with temporal attention. All accuracy and RT statistics for the combined experiment are reported in Table 1.

**Table 1.**
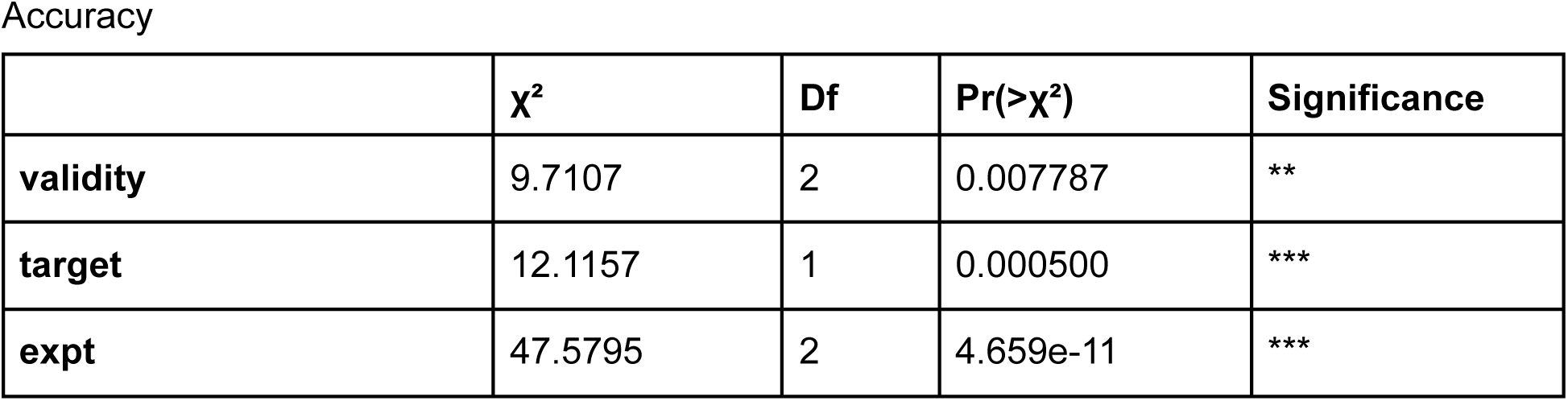
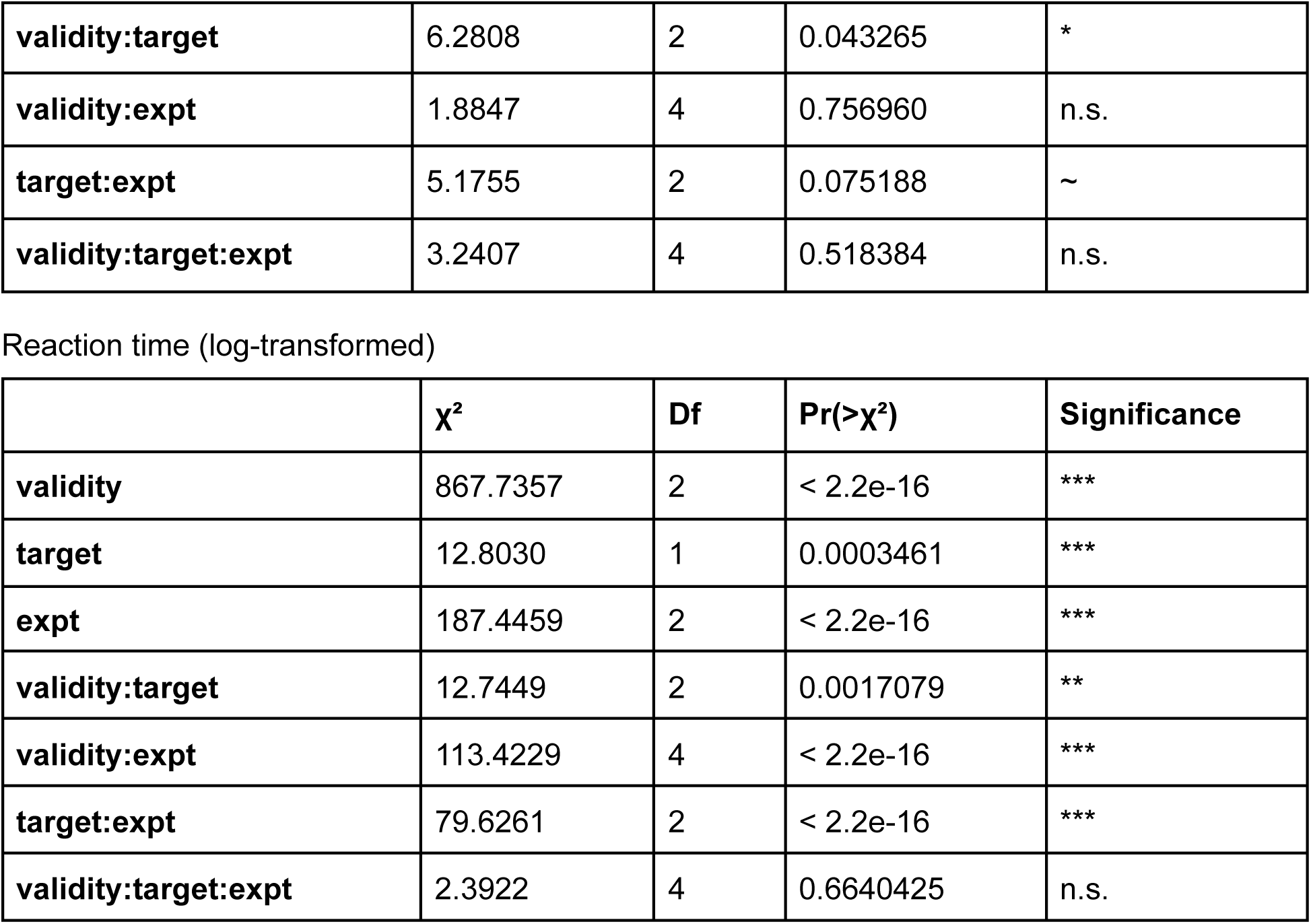
Combined experiment 1–3 statistics for accuracy and reaction time. ∼ p < 0.1, * p < 0.05, ** p < 0.01, *** p < 0.001.

Accuracy
| | $\chi^2$ | Df | Pr(>$\chi^2$) | Significance |
| --- | --- | --- | --- | --- |
| **validity** | 9.7107 | 2 | 0.007787 | ** |
| **target** | 12.1157 | 1 | 0.000500 | *** |
| **expt** | 47.5795 | 2 | 4.659e-11 | *** |
| **validity:target** | 6.2808 | 2 | 0.043265 | * |
| **validity:expt** | 1.8847 | 4 | 0.756960 | n.s. |
| **target:expt** | 5.1755 | 2 | 0.075188 | ~ |
| **validity:target:expt** | 3.2407 | 4 | 0.518384 | n.s. |

| | $\chi^2$ | Df | Pr(>$\chi^2$) | Significance |
| --- | --- | --- | --- | --- |
| **validity** | 867.7357 | 2 | < 2.2e-16 | *** |
| **target** | 12.8030 | 1 | 0.0003461 | *** |
| **expt** | 187.4459 | 2 | < 2.2e-16 | *** |
| **validity:target** | 12.7449 | 2 | 0.0017079 | ** |
| **validity:expt** | 113.4229 | 4 | < 2.2e-16 | *** |
| **target:expt** | 79.6261 | 2 | < 2.2e-16 | *** |
| **validity:target:expt** | 2.3922 | 4 | 0.6640425 | n.s. |

### Comparing auditory and visual temporal attention

The results so far demonstrate that temporal attention improved auditory perceptual discriminability only for T1. Further, we found auditory inattentional costs at T1 without corresponding benefits at T2, whereas the well-matched visual task (Denison et al., 2017) showed both benefits and costs that were mirrored across targets, with attentional benefits to one target accompanied by attentional costs to the other—indicating attentional tradeoffs. Because the current auditory temporal attention protocol was designed to closely match the visual one in task structure, timing, and the manipulation of temporal attention, we next quantitatively compared the current auditory data to the visual data from Denison et al., 2017. This contrast offers the opportunity to compare the effects of temporal attention across modalities.

## Methods

### Analysis

We first compared accuracy and RT averages for each participant and condition from Experiments 1, 2, and 3 to those from the visual temporal attention data of (Denison et al., 2017) Experiment 1 (n = 10). These datasets were modeled using LMMs. Our primary question of interest was whether there were any interactions of modality and validity, which would indicate a difference in attentional effects between modalities.

To visualize differences in attentional effects, we computed participant-level difference scores (valid – invalid for accuracy; invalid – valid for RTs). We then compared these scores between experiments using target-specific models. For auditory vs. visual comparisons, we used standard linear models (LMs), as the datasets did not share subjects and random effects could not be estimated. For completeness, we also directly compared the difference scores between the auditory Experiments 1 and 2 using linear mixed-effects models (LMMs) with subject-specific random intercepts, since these datasets included overlapping participants. Auditory experiment 3, however, did not share participants with any other experiment; therefore we also used standard linear models (LMs) when comparing it to the others.

## Results

### Attention improved perceptual accuracy in vision more than in audition

Temporal attention had more pronounced effects for visual discriminability than those of auditory discriminability (interaction between modality and validity, *χ²*(2) = 10.29, *p* = 0.006). This pattern was consistent for each target separately (T1: *χ²*(2) = 23.97, *p* = 6.22e-06; T2: *χ²*(2) = 9.98, *p* = 0.007). Furthermore, all paired-experiment target difference scores were significantly higher in vision than in audition (*F*(1, 18–19) ≥ 10.94, *p* ≤ 0.004 for all) other than T2 in Experiment 2, which was trending (*F*(1, 18) = 3.96, *p* = 0.062) (Denison et al., 2017) (**Figure 5a**).

**Figure 5.**
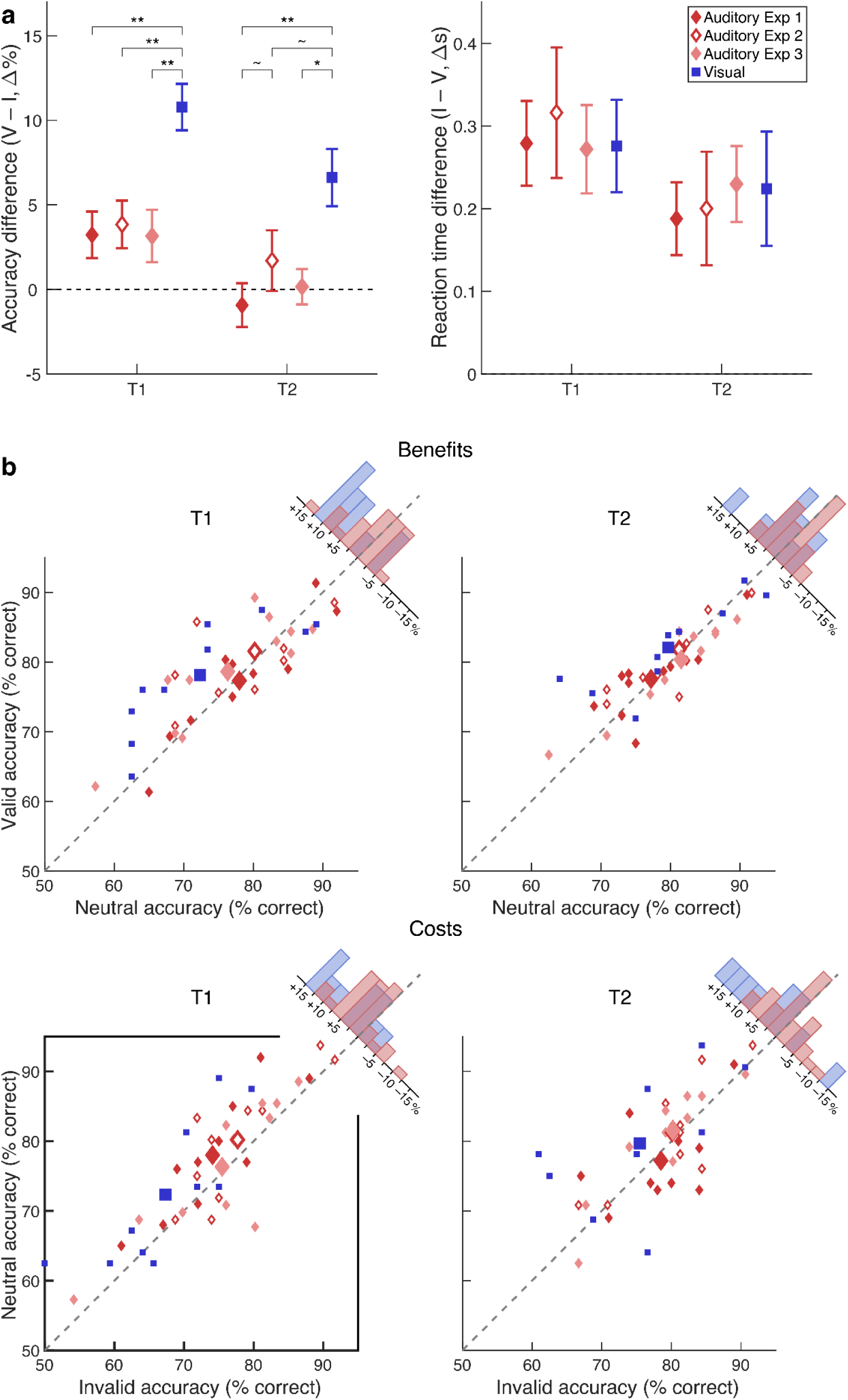
Comparing auditory and visual temporal attention. **(a)** Difference in accuracy and reaction time between valid and invalid conditions in Experiments 1–3, and previous visual temporal attention results, Experiment 1 from (Denison et al., 2017). **(b)** Comparison of individual subject and experiment average valid, neutral, and invalid accuracies for the same experiments showing attentional benefits (valid vs. neutral, top row) and costs (neutral vs. invalid, bottom row). Small dots represent individual subjects and large dots represent condition means for each experiment. The region above the diagonal unity line indicates attentional benefits (valid > neutral) or costs (neutral > invalid). The black box shows the only condition in which auditory temporal attention exhibits significant attentional effects (T1 costs), whereas visual temporal attention shows more symmetric benefits and costs across both targets. The diagonal histograms show the normalized distributions of subject-level attentional benefits and costs for the combined auditory experiments (red) and visual experiment (blue), obtained by projecting each subject’s scatterplot point onto the axis perpendicular to the unity line (i.e., the difference between the y- and x-axis conditions). Note this results in both positive benefits and positive costs being plotted to the left of the histogram zero point. Error bars represent ±1 SEM. n = 10 participants for each experiment. V = valid, N = neutral, I = invalid. ∼ p < 0.1, * p < 0.05, ** p < 0.01.

There was no significant interaction between modality and validity on RTs for the main model or either target submodel (*χ*²(2) ≤ 0.06, *p* ≥ 0.970 for all). Comparing the pairwise difference scores, RTs in each auditory task were indistinguishable from those in the visual tasks for each target (invalid – valid; *F*(1, 18–19) ≤ 0.20, *p* ≥ 0.663 for all) (**Figure 5a**).

Finally, difference scores were matched between any two auditory experiments, both in accuracy (valid – invalid; T1: *χ*²(1) or *F*(1, 19) ≤ 0.21, *p* ≥ 0.648 for all; T2: *χ*²(1) or *F*(1, 19) ≤ 3.17, *p* ≥ 0.075 for all) and in RT (invalid – valid; T1: *χ*²(1) or *F*(1, 19) ≤ 0.222, *p* ≥ 0.643 for all; T2: *χ*²(1) or *F*(1, 19) ≤ 1.12, *p* ≥ 0.290 for all) (Denison et al., 2017) (**Figure 5a**). There were no three-way interactions of modality, validity, and target for accuracy or RT (*χ²*(2) ≤ 0.27, *p* ≥ 0.872 for all). Overall, these results suggest that temporal attention leads to greater changes in accuracy in the visual domain than in the auditory domain.

Next, we compared the pattern of attentional benefits (valid vs. neutral) and costs (neutral vs. invalid) between the visual and auditory modalities by plotting each individual participant’s mean accuracy across targets and cue conditions (**Figure 5b**). In the visual experiment, most individual data points lie above the unity line for both targets, reflecting benefits and costs of temporal attention. In contrast, the auditory experiment only exhibited T1 costs, as shown by individual data points distributed above the unity line, whereas there was no indication of T1 benefits, T2 benefits, or T2 costs in audition (**Figure 5b**).

## Discussion

We investigated whether voluntary temporal attention can enhance auditory discrimination at attended time points, and whether such enhancement incurs performance costs at unattended time points. Across three experiments, we observed that temporal cueing improved auditory perception—specifically, frequency sweep direction discrimination—for the first target, T1, with no significant improvement for the second target, T2. Furthermore, the effect of temporal attention was independent of feature uncertainty. Finally, auditory perception was no better when one target was attended than when both were; its effects at T1 were associated with significant costs only. These findings indicate that, unlike for visual perception, temporal attention can enhance auditory perception without producing the reciprocal pattern of benefits and costs across time predicted by a limited-resource account. Indeed we found three converging lines of evidence against the idea that auditory processing resources are limited on this timescale: (1) attention did not affect both targets, (2) attention did not produce benefits at either target, (3) attention produced costs at T1 without reciprocal benefits at T2. Overall, the results suggest that attentional allocation in audition may operate under different temporal constraints than in vision.

### The absence of tradeoffs was specific to perceptual discrimination

While changes in attention did not exhibit a reciprocal pattern of benefits and costs in perceptual discriminability across time, the validity of the precue did improve the efficiency of response selection. Reaction times were fastest with valid and slowest with invalid precues for both targets. In this task, however, reaction time likely reflects the efficiency of response selection rather than perception per se, because the response cue only occurs after both targets have been presented. With a valid response cue, participants can therefore continue with any response plan developed during the trial (Correa, 2010), resulting in shorter reaction times, whereas with an invalid response cue, participants must switch any prepared response upon receiving the cue, resulting in longer reaction times. Further, participants were instructed to focus on accuracy and make an unspeeded judgment, with a delay of at least 500 ms enforced by having to wait for the response window. In Experiment 3, the response window started 1.1 s after the second stimulus onset, well after perceptual processing is expected to be complete (Donkin et al., 2014; Vangkilde et al., 2012). So although reaction time provides information about the speed of response selection following the response cue, it is unlikely to track changes in perceptual processing speed in this task protocol. Altogether, even though auditory temporal attention affected perceptual discriminability with costs at T1 only, the RT findings indicate that precue validity improved the efficiency of response selection across both targets, as expected, and confirm that participants followed the task instructions.

### Feature uncertainty did not modulate attentional effects

Although Experiment 2 reduced feature uncertainty by limiting the range of possible central frequencies compared to Experiment 1, the overall pattern of attentional effects remained unchanged: participants showed improved performance for T1, no significant attentional modulation for T2, and no evidence of tradeoffs. These results suggest that the benefits of temporal attention were robust to differences in feature predictability.

Feature uncertainty has been shown to impair discrimination and detection performance in both low- and high-level visual tasks (Irons et al., 2012; Kim et al., 2019; Mestry et al., 2017; Michel & Geisler, 2011; Pelli, 1985; Vogels et al., 1988). When the relevant stimulus dimension is uncertain or cued only after stimulus presentation, observers exhibit larger just-noticeable differences and lower sensitivity (Pelli, 1985; Vogels et al., 1988), likely reflecting increased demands on feature selection and sensory representation. Uncertainty also constrains attentional control settings, reducing the precision with which attention can be tuned to a relevant feature dimension (Irons et al., 2012). If temporal attention operates by modulating feature-tuned populations, we might predict that such modulation would be less effective with higher feature uncertainty, when more populations would need to be targeted by top-down attentional mechanisms. On the other hand, feature uncertainty about stimulus orientation increased the effects of visual temporal attention on orientation judgments (Palmieri & Carrasco, 2024). At least in vision, temporal attentional benefits seem to depend on the feature uncertainty of the task.

In our task, however, reducing uncertainty about the central frequency improved both discrimination thresholds and overall accuracy but did not alter the relative benefits of valid temporal cues—consistent with additive rather than interactive effects of these two factors. Therefore temporal attention may enhance auditory discrimination by modulating when processing resources are engaged without needing to target feature-specific neural populations.

### Auditory versus visual temporal attention

Temporal attention has been hypothesized to operate as a limited but recoverable resource in vision, where boosting perceptual discriminability at one time point incurs costs at another (Denison et al., 2021, 2024; Duyar et al., 2024). By adapting an established visual temporal attention paradigm (Denison et al., 2017) to the auditory domain, our study provided a between-modality comparison of temporal attention under tightly matched task conditions. Both modalities showed improved perceptual discriminability following valid temporal cues, consistent with the general capacity of voluntary attention to prioritize expected moments in time (Coull & Nobre, 1998; Nobre & van Ede, 2018). However, the magnitude and temporal profile of these benefits diverged: compared to visual attention, auditory attention yielded smaller effects restricted to T1.

Moreover, these attentional effects on T1 arose from costs on invalid trials without corresponding T2 benefits. This benefit-cost pattern is qualitatively different from the patterns found in visual studies. Visual temporal attention often shows reciprocal benefit-cost patterns: benefits in one target tend to be accompanied by costs in the other (Denison et al., 2017, 2021, 2024; Zhu et al., 2024). In contrast, the present auditory data showed only a T1 cost, with no T1 benefit and no reliable T2 benefit or cost. This pattern therefore differs from the reciprocal redistribution of perceptual resources expected under a limited-resource account. The different benefit-cost patterns for audition and vision may therefore suggest distinct attentional constraints for the two modalities.

Despite the difference in perceptual discriminability effects, RT patterns were strikingly consistent across modalities: valid cues sped responses, and invalid cues slowed them, with quantitatively similar changes in RT across modalities. This similarity suggests that both visual and auditory attention engage shared cognitive processes—such as temporal orienting or task preparation (Nobre & van Ede, 2018)—even if the consequences for perception differ across modalities.

Although the auditory and visual tasks were well-matched and task difficulty was comparable through staircasing, the larger effects of temporal attention in vision could reflect different factors. It may be that there is genuinely more effective temporal allocation for vision than for audition. Alternatively, given the impossibility of asking participants to discriminate identical stimulus features in auditory vs. visual tasks, it may be that attention affects orientation discrimination more than frequency discrimination for stimulus-specific reasons. However, the absence of reciprocal tradeoffs in audition—costs at T1 without corresponding benefits at T2—suggests that auditory temporal attention follows a different temporal profile, operating under distinct temporal or representational constraints from its visual counterpart. Although the generally weaker effects for audition may mean that there is an effect of temporal attention for T2 smaller than what we can measure, there was no hint of such an effect in the high-powered combined dataset.

Future studies will be needed to determine how temporal attention affects auditory perception across a range of stimuli and tasks, but the current study suggests qualitative differences in the allocation of temporal attention in audition and vision.

### Mechanisms underlying the absence of tradeoffs in auditory temporal attention

The pattern of behavioral accuracy we observed—attentional costs for T1 and no effect of attention for T2—suggests a hypothesis for the dynamics of attentional allocation in this task. When the precue is to T1 or neutral, attention is allocated to T1 and can also be allocated to an equal degree to T2, resulting in high performance for both targets. But when the precue is to T2, attention is not allocated until T2, resulting in costs for T1 on invalid trials. Such dynamics would arise in a system that followed the precue instructions to initiate attentional allocation (attend to T1, attend to T2, attend both) but had no temporal constraints limiting its ability to attend to both targets. We consider three mechanistic explanations for the apparent absence of such constraints at this timescale in audition.

First, auditory temporal attention and visual temporal attention might operate under different temporal constraints, or limits in processing ability over a certain timeframe. In vision these constraints operate over several hundred milliseconds (Denison, 2024; Denison et al., 2021), but our results raise the possibility that auditory constraints have a shorter duration, such that by 250 ms—the SOA used in our task—the system has already reset and can allocate resources to both T1 and T2 without interference. The dynamic normalization model of temporal attention posits that the source of temporal constraints is a limited availability of attentional gain across short time intervals (Denison et al., 2021). Within this framework, we might hypothesize that attentional gain recovers more rapidly for audition than for vision.

A second possibility is that the auditory system supports a more sustained (Lange et al., 2003) gain mechanism—one that can modulate perception over a broader time window without depleting resources from nearby time points. This possibility would entail a fundamental difference in how attention is allocated across time in audition versus vision.

Third, the auditory system’s natural tendency to group sequential inputs into perceptual streams may reduce the need for attentional reallocation to the second target. Even though our task involved discrete stimuli, T1 and T2, streaming may have enabled integration across the two stimuli, allowing modulation directed at one time point to spread across the entire stream (Noyce et al., 2023; Shinn-Cunningham, 2008). Consistent with this possibility, auditory stream formation depends on sustained attention and can reset when attention shifts (Cusack et al., 2004). Because our task design lacked strong perceptual boundaries between targets, attention may have spread over a single ongoing stream, enabling sustained modulation across both targets and mitigating tradeoffs or competition between them.

Taken together, these findings suggest that the absence of attentional tradeoffs in our study may reflect a combination of factors: (1) faster recovery of attentional gain in audition, (2) more sustained auditory attentional modulation over time, and (3) temporal grouping of stimuli into coherent auditory streams. These mechanisms are not mutually exclusive, and any of them could allow attention to remain elevated across successive targets without incurring costs—at least under the current task conditions and timing. As auditory perception emphasizes the continuity and integration of information over time (Noyce et al., 2023; Shinn-Cunningham, 2008), these findings highlight the need to consider how temporal attention is shaped by the representational and ecological demands of each sensory modality.

### Future directions

Our study provides an initial demonstration that temporal cueing can enhance auditory discrimination without incurring tradeoffs. To build on this foundation, future work could explore a broader range of stimulus onset asynchronies (SOAs) to test whether auditory attention shows different dynamics at shorter or longer intervals. In addition, we focused on a single auditory task—sweep direction discrimination—which offers a well-controlled test of perceptual discriminability but may not capture the full range of auditory processing demands. Extending this protocol to other auditory judgments would clarify the generalizability of these effects. Finally, introducing more complex auditory scenes with competing streams or distractors could more directly probe the capacity limits of auditory temporal attention under realistic listening conditions.

## Conclusion

In summary, temporal cueing enhances auditory discrimination without clear evidence of tradeoffs in perceptual discriminability, contrasting with previously documented visual tradeoffs. These findings suggest that while temporal attention improves perception in both modalities, the underlying dynamics and temporal constraints may differ. Auditory temporal attention may be able to be more sustained across successive stimuli, reflecting the modality’s temporal demands and ecological function.

## Acknowledgments

This research was supported by NIH R01EY037358 and Boston University startup funding to R.N.D. and Undergraduate Research Opportunities Program funding to J.W. We thank the members of the Denison Lab and Stephanie Badde for helpful discussions of the project.

## Resource availability

The data and all experiment and analysis code will be made publicly available upon publication.

## Author contributions

Conceptualization: J.W., R.N.D. Methodology: J.W., C.C., R.N.D. Data collection: J.W. Data analysis: J.W. Writing: J.W., R.N.D. Review & Editing: J.W., C.C., R.N.D.

